# cit_gsa: gene set analysis for scRNA-seq with covariate adjustment

**DOI:** 10.64898/2026.10.13.758831

**Authors:** Sara Fallet, Véronique Godot, Pierre Neuvial, Boris P. Hejblum

## Abstract

**Summary:** Single-cell RNA-seq enables gene expression analysis at single-cell resolution. Differential Expression Analysis (DEA) performed at the gene set level for better power and interpretation, identifies gene sets associated with a factor of interest. Existing single-cell DEA methods rely on two-step approaches, a gene-level analysis followed by an enrichment step, are often limited to two-group comparisons and do not support covariate adjustment. We introduce `cit_gsa`, a novel one-step self-contained gene set DEA method for Single-cell RNA-seq that handles more than two conditions and allows covariate adjustment. A benchmark of `cit_gsa` against state-of-the-art methods shows that our method has better performance in complex scenarios, and real-data analysis confirms that covariate adjustment improves biological interpretation.

**Availability and implementation:** `cit_gsa` is implemented in an R package available at https://CRAN.R-project.org/package=citcdf. The data and R scripts are available at https://doi.org/10.5281/zenodo.23264404 for reproducibility.

## Introduction

Single-cell RNA-seq (scRNA-seq) allows the study of gene expression at the level of individual cells. This provides a finer and more precise resolution than bulk RNA sequencing, where only average gene expression across cells is available. However, scRNA-seq data are high-dimensional count data with numerous dropout events, and also exhibit dependencies between cells as well as correlation between genes.

Differential Expression Analysis (DEA) allows to study the variation of gene expression across experimental conditions, e.g. different vaccine arms. To model this data, count distributions such as the negative binomial or zero-inflated negative binomial are commonly used to account for overdispersion and dropout events. Many methods have been developed, such as `zingeR` (Van den Berge et al., 2018), `MAST` (Finak et al., 2015) or the Wilcoxon test. In order to avoid strong distributional assumptions, other non-parametric methods have been developed, such as `citcdf` (Gauthier et al., 2021) or `distinct` (Tiberi et al., 2023).

In practice, it is biologically more interesting to analyse gene sets instead of individual genes (Subramanian et al., 2005). Gene sets are groups of genes known to be linked to the same biological function. Studying gene sets allows us to gain statistical power in the analysis and to obtain more interpretable results. Gene sets can come from general databases based on prior biological knowledge, such as Gene Ontology (Ashburner et al., 2000), KEGG (Kanehisa and Goto, 2000), or MSigDB (Liberzon et al., 2011); or domain-specific databases built from co-expression observations such as the Blood Transcription Modules (Li et al., 2014) or BloodGen3 (Chaussabel et al., 2008) in immunology.

Gene set methods for differential analysis can be split into two categories according to the null hypothesis they test: self-contained and competitive methods (Goeman and Bühlmann, 2007). Self-contained methods test the null hypothesis that no gene in the gene set of interest is differentially expressed, e.g. `globaltest` (Goeman et al., 2004) originally developed for bulk data, or `SCPA` (Bibby et al., 2022) specifically developed for single-cell data. Competitive methods test the null hypothesis that the genes in the gene set of interest are not more differentially expressed than the other genes included in the study, e.g. `GSEA` (Subramanian et al., 2005) and the hypergeometric test (Rivals et al., 2007) are the most popular method originally developed for bulk data, and `TWO-SIGMA-G` (Van Buren et al., 2022) or `iDEA` (Ma et al., 2020) specifically tailored for single-cell data. Several comparisons have been made between self-contained and competitive methods (Maciejewski, 2014; Mathur et al., 2018). Goeman and Bühlmann (2007) state that self-contained methods should be used, as they provide statistically more interpretable results.

### Contribution

We introduce `cit_gsa` , a self-contained gene set analysis method allowing the comparison of more than two conditions, including continuous variables, while adjusting for covariates. Unlike existing single-cell methods, our method combines gene-level DEA and gene set enrichment in a single step. After describing `cit_gsa` , we perform a numerical comparison to competitors on simulated data. Then, we apply `cit_gsa` to Dendritic Cells data to demonstrate that the ability to adjust for covariates can influence the results of the analysis and provide additional biological insights.

## Method

Consider a gene set of size *s* (indexed by [*s*] := 1, … *s*) and denote by ***Y*** the *s*-dimensional vector of gene expressions within this gene set. Let ***X*** be an experimental factor of interest, such as a disease status or a vaccine arm. Note that ***X*** may be discrete or continuous. We also assume to be given some covariates ***Z***, such as the cell type. Both ***X*** and ***Z*** can be multivariate. Our aim is to test the conditional independence between ***Y*** and ***X*** given ***Z***, based on *n* observations corresponding to *n* cells. Formally, the null hypothesis for our test is:

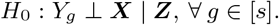

We introduce the `cit_gsa` method, a gene set-level extension of the `citcdf` method of Gauthier et al. (2021). `citcdf` performs a conditional independence test of a single gene-level null hypothesis: *H*_0*g*_ : *Y*_*g*_ ⊥ ***X*** | ***Z***. The null hypothesis of our test can be equivalently rewritten based on conditional cumulative distribution functions (cdfs):

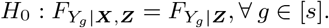

Here, 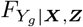 is the conditional cdf of the expression of gene *g* given both the factor and covariates, and 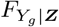 is the conditional cdf of the expression of gene *g* given the covariate only. This conditional cdf can be expressed as the conditional expectation of a binary variable: 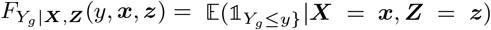. Without loss of generality, and for the sake of keeping notations light-weight, we consider *X* and *Z* uni-dimensional in the remainder but the method and its implementation allows to take into account several variables of interest *X* and covariates *Z* (see Supplementary Section S1). Our goal is to test *H*_0_ based on *n* independent, identically distributed observations of ***Y*** , *X* and *Z*, denoted by ((*Y*_*ig*_)_*g*∈[*s*]_, *X*_*i*_, *Z*_*i*_)_*i*∈[*n*]_, each observation corresponding to one cell. Following Gauthier et al. (2021), to do so we choose a sequence of *p* regular and ordered thresholds *ω*_*j*_ where *j* ∈ [*p*], and estimate the above expectations using a sequence of *p* logistic regressions:

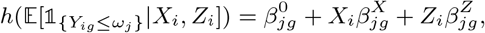

If the factor of interest *X* has no link with the gene expression *Y*_*g*_, then the vector 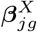 will be equal to 0. In practice (see Supplementary Section S1.1 for details), we test the hypothesis: “*H*_0_ : ***β***^*X*^ = 0” where 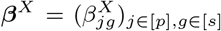. The associated score test statistic aggregates the corresponding estimators: 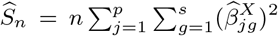. In Supplementary Section S1 we obtain a *p*-value for *H*_0_ by comparing the observed test statistic 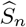 with its asymptotic distribution. The latter is shown to be a mixture of 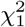 random variables: 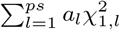, where the *a*_*l*_ are the eigenvalues of the variance-covariance matrix Σ of the ***β***^*X*^ . As we are in gene set analysis, Σ involves cross terms between the genes of the gene sets.

## Results

### Simulations

A natural competitor for `cit_gsa` is `globaltest` (Goeman et al., 2004), a self-contained method that uses a regression model to test whether the mean gene expression within gene sets is associated with the variable of interest *X*, while accounting for covariates ***Z***. `cit_gsa` can also be compared to the most widely used gene set analysis approaches, which are enrichment methods with a competitive test: the hypergeometric test and `GSEA` . The hypergeometric test (Rivals et al., 2007) assesses whether a gene set is overrepresented relative to a list of genes called differentially expressed (DE). `GSEA` (Subramanian et al., 2005) ranks genes by gene-level statistic and computes an enrichment score whose significance can be based on gene or label permutations. We have used gene permutations as explained in Supplementary Section S2.1. As they both require gene-level statistics as an input, we use the *p*-values from the gene-level `citcdf` testing. Simulations have been done in `R` v4.5.1 with the `citcdf` , `globaltest` , and `fgsea` packages.

#### Framework and metrics

We compared two DE scenarios: i) the classical approach involving a difference in mean between the conditions (DMean), and ii) a difference in modality (DMod), i.e. that the number of modes in the distribution of truly DE genes differs across conditions, which is relevant in single-cell RNA-seq (Korthauer et al., 2016). We generated *m* = 1000 genes and *n* = 200 cells and we tested one gene set of 100 genes. We varied the proportion of genes under the alternative (i.e. truly DE genes) in the gene set between 0 and 1, while the total number of truly DE genes in the expression matrix always remained 500. The number of permutations was set to 1000.

The covariate *Z* ∈ ℝ^*n*^ and the variable of interest *X* ∈ ℝ^*n*^ were generated according to Gaussian distributions. The gene expression matrix *Y* ∈ ℝ^*n*×*m*^ is defined as: *Y* = signal + *ϵ*, where *ϵ* is an equi-correlated Gaussian noise matrix of parameter *ρ* ∈ {0, 0.4, 0.8} which is the correlation between any pair of genes. The parameter `signal` depends of the scenario DMean/DMod, as described in Supplementary Section S2.2.

To compare the results of each method, we report the True Discovery Rate (TDR), which corresponds to the average proportion of *p*-values below 0.05, across 500 replications of this experiment.

#### Results

Figure 1 represents TDR as a function of the proportion of truly DE genes in a gene set of size 100 for the two scenarios DMean and DMod. The correlation is *ρ* = 0.4. Self-contained methods are found to be more powerful than competitive ones, in line with the fact that the null hypothesis of self-contained methods is more restrictive. While both self-contained methods have similar power in the DMean scenario, `cit_gsa` is more powerful than `globaltest` in the DMod scenario. Indeed, the latter is based on a regression model for the conditional mean of the gene expression, which cannot detect changes in modality. In contrast, `cit_gsa` is based on the comparison of conditional distribution functions, which is more powerful in this context. Similar conclusions hold for other levels of dependence between genes (*ρ* ∈ {0, 0.8}), see Supplementary Section S2.3.

**Figure 1.**
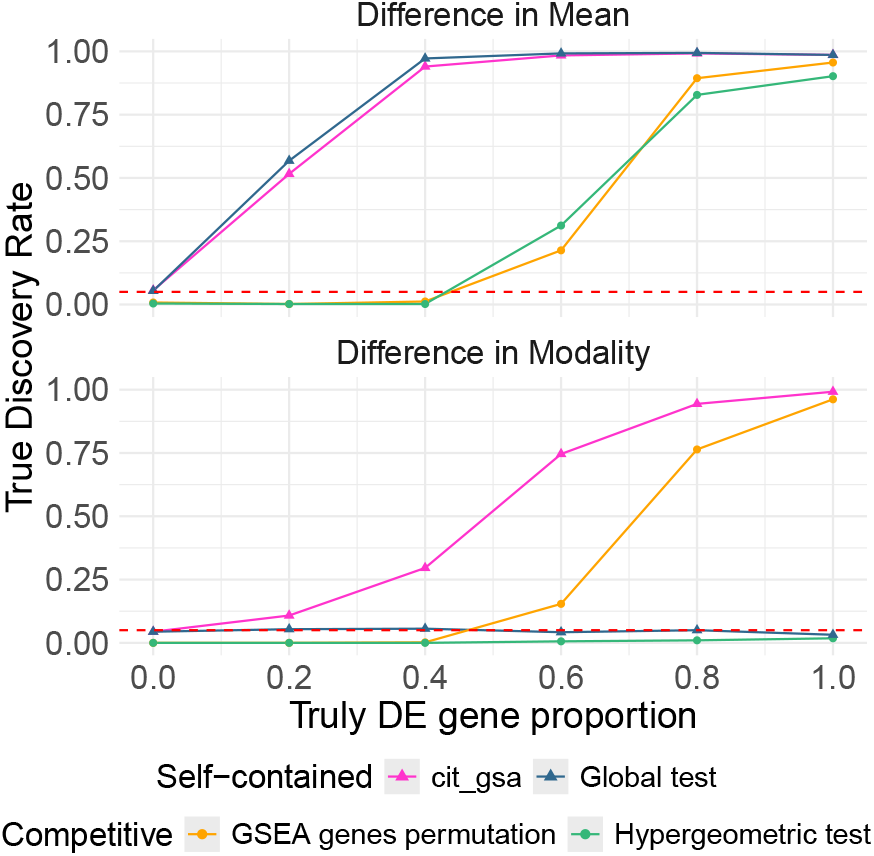
True Discovery Rate as a function of the proportion of truly DE genes in a gene set of size 100, for two DE scenarios: DMean (top) and DMod (bottom) with *ρ* = 0.4.

### Dendritic Cells data analysis

#### Dataset

Dendritic Cells (DCs) are immune cells responsible for antigen-presentation, i.e. capturing, processing, and presenting antigens to T cells to initiate the adaptive response of the immune system when faced with a pathogen. Villani et al. (2017) isolated DCs from Peripheral Blood Mononuclear Cells (PBMCs) in healthy donor blood, and measured gene expression through single-cell RNA-seq. Villani et al. (2017) documented distinct DC subpopulations with each having their own specific transcriptional signatures (Supplementary Figure S3). These subpopulations and their respective signatures were identified through graph-based clustering followed by marker-genes analysis comparing each cell cluster to the others.

#### Scientific question

Quality control processing and normalization with log-transformed transcripts per million (log(*TPM* + 1)) of the data from Villani et al. (2017) yields 539 single cells observed along which 17,691 genes. We performed a differential gene set analysis to identify which of the Blood Transcription Modules (BTM) from Li et al. (2014) are significantly associated with the expression of CD83, known for being a marker of DCs activation (Zhou and Tedder (1995); Lechmann et al. (2002)). Because CD83 expression varies across DC subsets (Supplementary Figure S4) and since `cit_gsa` allows for covariate adjustment, we included DCs subpopulation as a covariate in our differential gene set analysis. This ensures that any association found between CD83 expression and gene set expression reflects a biological link to activation, rather than differences between DC subsets. Supplementary Section S3.3 provides a similar analysis with IDO1, a gene associated with immune tolerance of DCs.

#### Results

Each dot in Figure 2 corresponds to one of the 259 annotated BTM from Li et al. (2014). The *x*-axis shows the *conditional p*-values obtained with `cit_gsa` when adjusting for DC subsets, whereas the *y*-axis shows the corresponding *marginal p*-values obtained with no such adjustment. The *p*-values are shown in logarithmic scale, after performing a False Discovery Rate (FDR) multiple testing correction (Benjamini and Hochberg, 1995).

**Figure 2.**
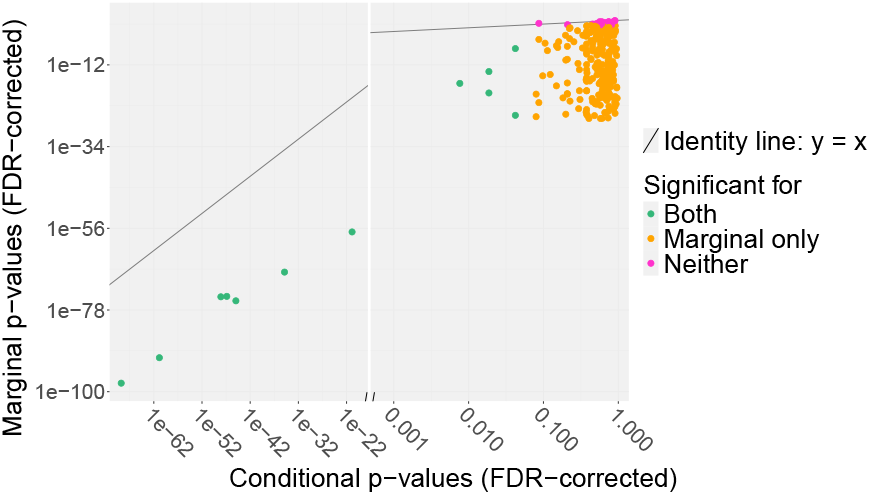
Gene set analysis results for the association between CD83 expression, comparing for each of the 259 BTM gene sets of Li et al. (2014) the conditional *p*-value after DC subset adjustment (*x*-axis) with the marginal *p*-value with no such adjustment (*y*-axis). The FDR-corrected *p*-values are shown on a logarithmic scale, with a break in the *x*-axis. The solid line represents the identity line.

Among the 259 annotated gene sets, 237 (shown as orange and green dots in Figure 2) are significantly associated with CD83 when no covariate adjustment is performed. After adjusting for DC subsets, only 12 gene sets (green dots) remain significant. Their names are provided in Supplementary Table S1. This result suggests that many of the associations observed without adjustment are driven by differences between DC subsets rather than by CD83 expression itself.

## Conclusion

We introduce `cit_gsa` , a new self-contained method for differential gene set analysis of single-cell RNA-seq data. Unlike many existing approaches which are limited to two-group comparisons, our method can handle multiple variables of interest. In addition, it allows adjustment for covariates, reducing the impact of potential confounding factors on the results and enabling a more accurate interpretation of the associations between gene sets and the variable of interest.

The simulation study shows that `cit_gsa` is more powerful than competitive approaches. It can also detect more complex differential patterns than existing self-contained methods, such as differences in the distribution of gene expression, a feature relevant to single-cell RNA-seq data. As a general non-parametric approach, `cit_gsa` is not limited to single-cell data and can also be applied to other types of transcriptomic data such as bulk RNA-seq.

For now, `cit_gsa` does not account for the correlation between cells that come from the same biological sample. Some methods have recently been developed, specifically for single cell, to address this issue in gene set analysis. For example, `TWO-SIGMA-G` incorporates a random effect into a gene-level mixed model to account for multi-sample correlation before performing gene set enrichment analysis. Extending our framework to account for multi-sample data while preserving its self-contained testing strategy is a direction for future work.

## Supplementary material

Supplementary materials are available at Bioinformatics online.

## Conflicts of interest

The authors declare that they have no competing interests.

## Funding

This work is supported by Agence Nationale de la Recherche under the France 2030 program (PEPR Santé Numérique AI4scMED ANR-22-PESN-0002) and under France’s PIA 3 scheme through the University of Bordeaux’s Digital Public Health Graduate School (Investments for the Future-Project reference: 17-EURE-0019). Computer time for this study was provided by the computing facilities MCIA (Mésocentre de Calcul Intensif Aquitain) of the Université de Bordeaux and of the Université de Pau et des Pays de l’Adour.

## Data availability

The R package `citcdf` available on CRAN implements `cit_gsa` . All codes and data to reproduce the results of this article are publicly accessible from Zenodo at https://doi.org/10.5281/zenodo.23264404.

## Notes

### Competing Interest Statement

The authors have declared no competing interest.

