## Supplementary Materials for "cit_gsa: gene set analysis for scRNA-seq with covariate adjustment"

October 13, 2026

[1]Univ. Bordeaux, INSERM, INRIA, Bordeaux Population Health, SISTM team, U1219, F-33000 Bordeaux, France.
[2]Institut de Mathématiques de Toulouse, UMR 5219, Université de Toulouse, CNRS, UPS, F-31062 Toulouse Cedex 9.
[3]Vaccine Research Institute, Créteil F-94010, France.
[4]INSERM-U955, team Accelerates, Créteil F-94010, France.
[5]Faculté de Santé, Université Paris-Est, Créteil, France.
[6]IHU SEPSIS - PROMETHEUS Comprehensive Sepsis Center, Paris-Saclay University, Saclay, France.

### S1 Complements on the cit_gsa method

#### S1.1 Test hypothesis

Our objective is to test the null hypothesis:

$$H_0 : F_{Y_g|\boldsymbol{X},\boldsymbol{Z}} = F_{Y_g|\boldsymbol{Z}}, \forall\, g \in [s].$$

As explained in the main paper, following Gauthier et al. (2021), we approximate the conditional cumulative distribution function by using a sequence of $p$ thresholds $(\omega_j)_{j\in[p]}$ and we estimate the conditional cumulative distribution function by using a regression model for each threshold. Our working model is

$$h(\mathbb{E}[\mathbb{1}_{\{Y_{ig}\leq\omega_j\}}|\boldsymbol{X}_i, \boldsymbol{Z}_i]) = \beta^0_{jg} + \boldsymbol{X}_i^\intercal \boldsymbol{\beta}^{\boldsymbol{X}}_{jg} + \boldsymbol{Z}_i^\intercal \boldsymbol{\beta}^{\boldsymbol{Z}}_{jg} \tag{S1.1}$$

with $\boldsymbol{X}_i \in \mathbb{R}^K$, $\boldsymbol{Z}_i \in \mathbb{R}^q$, $i \in [n]$. Note that contrary to the main manuscript where we limited ourselves to the simpler case where the variable of interest $X$ is scalar (for the sake of clarity), we consider here the more general case of multidimensional $\boldsymbol{X}$ and $\boldsymbol{Z}$. In practice, we do not test $H_0$ but

$$H_0' : \boldsymbol{\beta}^{\boldsymbol{X}} = 0,$$

where $\boldsymbol{\beta}^{\boldsymbol{X}} = (\beta^{\boldsymbol{X}}_{jg})_{j\in[p],g\in[s]}$. As long as the linear dependency on $\boldsymbol{Z}$ is correctly specified, $H_0$ implies $H_0'$, so we use $H_0'$ as an approximation of $H_0$. In the case of a large number of thresholds $p$, it provides a close approximation of $H_0$.

#### S1.2 Estimation and test statistic

The above working model (S1.1) can be written as:

$$h(\mathbb{E}[D_{ij}^g|\boldsymbol{W}_i]) = \boldsymbol{W}_i^\intercal \boldsymbol{\beta}_{jg} \tag{S1.2}$$

for the $i$-th observation, $j$-th gene expression threshold and $g$-th gene, where $\boldsymbol{W}_i = (1, \boldsymbol{X}_i, \boldsymbol{Z}_i)^\intercal$ gathers the covariates for observation $i$ and $D_{ij}^g = \mathbb{1}_{\{Y_{ig} \leq \omega_j\}}$ is a binary variable indicating whether the expression of gene $g$ in observation $i$ is at or below the threshold $\omega_j$.

For computational efficiency reasons, and following Gauthier et al. (2021), we use the identity link function ($h(y) = y$) and estimate $\boldsymbol{\beta}_{jg}$ with Ordinary Least Squares (OLS), as an approximation to logistic regression. Letting $\boldsymbol{D}_{\cdot j}^g$ denote the $j$-th column of the binary matrix $\boldsymbol{D}^g = (D_{ij}^g)_{i\in[n], j\in[p]}$, the OLS estimator of $\boldsymbol{\beta}_{jg}$ is

$$\widehat{\boldsymbol{\beta}}_{jg} = (\boldsymbol{W}^\intercal \boldsymbol{W})^{-1} \boldsymbol{W}^\intercal \boldsymbol{D}_{\cdot j}^g,$$

with $\boldsymbol{W}$ the $n \times (1 + K + q)$ matrix whose $i$-th row is $\boldsymbol{W}_i^\intercal$.

Stacking the $\boldsymbol{X}$-coordinates of all the estimated coefficients into $\widehat{\boldsymbol{\beta}}^{\boldsymbol{X}} = (\widehat{\beta}_{jg}^{X^k})_{k\in[K], j\in[p], g\in[s]} \in \mathbb{R}^L$ with $L = Kps$, we define the observed gene-set test statistic as

$$\widehat{S}_n \;=\; n \left\|\widehat{\boldsymbol{\beta}}^{\boldsymbol{X}}\right\|^2 \;=\; n \sum_{j=1}^{p} \sum_{g=1}^{s} \sum_{k=1}^{K} \big(\widehat{\beta}_{jg}^{X^k}\big)^2. \tag{S1.3}$$

This statistic corresponds to the sum of the gene-level statistics introduced by Gauthier et al. (2021) over all genes in the considered gene set.

#### S1.3 Asymptotic null distribution

The population counterpart of $\widehat{\boldsymbol{\beta}}_{jg}$ is $\boldsymbol{\beta}_{jg}^* = \boldsymbol{\Psi}^{-1}\mathbb{E}[\boldsymbol{W}_i D_{ij}^g]$, with $\boldsymbol{\Psi} = \mathbb{E}[\boldsymbol{W}_i \boldsymbol{W}_i^\intercal]$. We denote the corresponding residuals by $\varepsilon_{ijg} = D_{ij}^g - \boldsymbol{W}_i^\intercal \boldsymbol{\beta}_{jg}^*$ and their vector over observations by $\boldsymbol{\varepsilon}_{\cdot jg} = \boldsymbol{D}_{\cdot j}^g - \boldsymbol{W}\boldsymbol{\beta}_{jg}^*$. So we can rewrite $\widehat{\boldsymbol{\beta}}_{jg}$ as:

$$\begin{aligned}
\widehat{\boldsymbol{\beta}}_{jg} &= (\boldsymbol{W}^\intercal \boldsymbol{W})^{-1} \boldsymbol{W}^\intercal \boldsymbol{D}_{\cdot j}^g \\
&= (\boldsymbol{W}^\intercal \boldsymbol{W})^{-1} \boldsymbol{W}^\intercal (\boldsymbol{W}\boldsymbol{\beta}_{jg}^* + \boldsymbol{D}_{\cdot j}^g - \boldsymbol{W}\boldsymbol{\beta}_{jg}^*) \\
&= \boldsymbol{\beta}_{jg}^* + (\boldsymbol{W}^\intercal \boldsymbol{W})^{-1} \boldsymbol{W}^\intercal \boldsymbol{\varepsilon}_{\cdot jg}
\end{aligned}$$

so that

$$\begin{aligned}
\sqrt{n}\,(\widehat{\boldsymbol{\beta}}_{jg} - \boldsymbol{\beta}_{jg}^*) &= (n^{-1}\boldsymbol{W}^\intercal \boldsymbol{W})^{-1} \frac{1}{\sqrt{n}} \sum_{i=1}^{n} \boldsymbol{W}_i \varepsilon_{ijg} \\
&= \boldsymbol{\Psi}^{-1} \frac{1}{\sqrt{n}} \sum_{i=1}^{n} \boldsymbol{W}_i \varepsilon_{ijg} + \Big\{(n^{-1}\boldsymbol{W}^\intercal \boldsymbol{W})^{-1} - \boldsymbol{\Psi}^{-1}\Big\} \frac{1}{\sqrt{n}} \sum_{i=1}^{n} \boldsymbol{W}_i \varepsilon_{ijg}.
\end{aligned} \tag{S1.4}$$

By the law of large numbers, $n^{-1}\boldsymbol{W}^\intercal \boldsymbol{W} \xrightarrow{P} \boldsymbol{\Psi}$, hence assuming that $\boldsymbol{\Psi}$ is invertible, then by the continuous mapping theorem $\boldsymbol{W}^\intercal \boldsymbol{W}$ is itself invertible for large $n$ and $(n^{-1}\boldsymbol{W}^\intercal \boldsymbol{W})^{-1} \xrightarrow{P} \boldsymbol{\Psi}^{-1}$. Besides, $\frac{1}{\sqrt{n}} \sum_{i=1}^{n} \boldsymbol{W}_i \varepsilon_{ijg}$ is a normalized sum of i.i.d. terms, centered by construction ($\mathbb{E}[\boldsymbol{W}_i \varepsilon_{ijg}] = \mathbb{E}[\boldsymbol{W}_i D_{ij}^g] - \boldsymbol{\Psi}\boldsymbol{\beta}_{jg}^* = \boldsymbol{0}$, even if the working model is misspecified), and is therefore bounded in probability by the Central Limit

Theorem. The second term in (S1.4) is thus $o_P(1)\,O_P(1) = o_P(1)$. Keeping only the coordinates corresponding to $\boldsymbol{X}$, this becomes:

$$\sqrt{n}\,(\widehat{\boldsymbol{\beta}}^{\boldsymbol{X}}_{jg} - \boldsymbol{\beta}^{*\boldsymbol{X}}_{jg}) = \frac{1}{\sqrt{n}}\sum_{i=1}^{n}\boldsymbol{\gamma}_i\,\varepsilon_{ijg} + o_P(1), \tag{S1.5}$$

where $\boldsymbol{\gamma}_i = (\gamma_i^1, \ldots, \gamma_i^K)^\intercal$ gathers the $K$ coordinates of $\boldsymbol{\Psi}^{-1}\boldsymbol{W}_i$ corresponding to $\boldsymbol{X}$: $\gamma_i^k$ is the product of the row of $\boldsymbol{\Psi}^{-1}$ corresponding to $X^k$ with $\boldsymbol{W}_i$.

Stacking (S1.5) over all thresholds $j$ and genes $g$, the $L$-dimensional vectors $(\boldsymbol{\gamma}_i\varepsilon_{ijg})_{j\in[p],g\in[s]}$ are i.i.d. across the $n$ observations (indexed by $i$) and centered (as sub-vectors of $\boldsymbol{\Psi}^{-1}\boldsymbol{W}_i\varepsilon_{ijg}$ whose expectation is $\boldsymbol{0}$). The multivariate Central Limit Theorem then gives:

$$\sqrt{n}\,(\widehat{\boldsymbol{\beta}}^{\boldsymbol{X}} - \boldsymbol{\beta}^{*\boldsymbol{X}}) \xrightarrow{d} \mathcal{N}_L(\boldsymbol{0}, \boldsymbol{\Sigma}). \tag{S1.6}$$

$\boldsymbol{\Sigma}$ is the covariance matrix of a single term of the sum over $i$ in (S1.5), i.e. of the vector $(\gamma_i^k\varepsilon_{ijg})_{k\in[K],\,j\in[p],\,g\in[s]}$: its $k, j, g$ and $k', j', g'$ entry is $\mathrm{Cov}(\gamma_i^k\varepsilon_{ijg}, \gamma_i^{k'}\varepsilon_{ij'g'})$, which reduces to the second moment $\mathbb{E}[\gamma_i^k\gamma_i^{k'}\varepsilon_{ijg}\varepsilon_{ij'g'}]$ since these are centered. It captures the correlations between the thresholds for the same gene, and between the genes of the set.

Finally, under $H_0'$ (as long as the linear dependency on $\boldsymbol{Z}$ is correctly specified in (S1.1)), $\boldsymbol{\beta}^{*\boldsymbol{X}}_{jg} = \boldsymbol{0}$, so by (S1.6) and the continuous mapping theorem once again, we obtain the convergence of $\widehat{S}_n = \left\|\sqrt{n}\,\widehat{\boldsymbol{\beta}}^{\boldsymbol{X}}\right\|^2$ to the squared norm of a random vector distributed as $\mathcal{N}_L(\boldsymbol{0}, \boldsymbol{\Sigma})$. Since the squared norm is invariant under rotation, expressing this vector in the orthonormal basis of eigenvectors of $\boldsymbol{\Sigma}$ turns it into a sum of $L$ independent squared centered Gaussian variables, whose variances are the eigenvalues of $\boldsymbol{\Sigma}$. Thus, the asymptotic distribution of $\widehat{S}_n$ under the null hypothesis is a mixture of $L$ independent $\chi^2_1$ distributions whose weights $a_l$ are the eigenvalues of $\boldsymbol{\Sigma}$:

$$\widehat{S}_n \xrightarrow[n\to\infty]{d} \sum_{l=1}^{L} a_l\,\chi^2_{1,l}. \tag{S1.7}$$

Throughout, the thresholds $\omega_j$ are treated as fixed. In practice they are chosen from the data (e.g. along a regular grid between the observed extremes of each gene). In any case, it does not affect the null distribution in (S1.7) since $\boldsymbol{\beta}^{*\boldsymbol{X}}_{jg} = \boldsymbol{0}$ for every $\omega_j$ under $H_0'$.

#### S1.4 Estimation of $\Sigma$ and p-value computation

$\boldsymbol{\Sigma}$ is estimated by replacing $\mathbb{E}[\gamma_i^k\gamma_i^{k'}\varepsilon_{ijg}\varepsilon_{ij'g'}]$ by an empirical mean, and by plugging in $n^{-1}\boldsymbol{W}^\intercal\boldsymbol{W}$ for $\boldsymbol{\Psi}$ in the estimation of $\gamma$ as well as the OLS residuals for $\varepsilon_{ijg}$. The $k, j, g$ and $k', j', g'$ entry of $\widehat{\boldsymbol{\Sigma}}$ is thus computed as follows, the between-gene terms ($g \neq g'$) being absent from Gauthier et al. (2021) who only consider one gene at a time:

$$\frac{1}{n}\sum_{i=1}^{n}\widehat{\gamma}_i^k\widehat{\gamma}_i^{k'}\,\widehat{\varepsilon}_{ijg}\,\widehat{\varepsilon}_{ij'g'}, \qquad \text{with} \quad \widehat{\varepsilon}_{ijg} = D_{ij}^g - \boldsymbol{W}_i^\intercal\widehat{\boldsymbol{\beta}}_{jg}, \tag{S1.8}$$

and where $\widehat{\boldsymbol{\gamma}}_i$ gathers the coordinates of $(n^{-1}\boldsymbol{W}^\intercal\boldsymbol{W})^{-1}\boldsymbol{W}_i$ corresponding to $\boldsymbol{X}$.

This corresponds to the multivariate version of the heteroskedasticity-robust sandwich estimator from White (1980). This estimator is consistent without needing to assume that the working model is correct (nor that the variance of $D_{ij}^g$ is constant across observations). Therefore, $\widehat{\mathbf{\Sigma}} \xrightarrow{P} \mathbf{\Sigma}$, and its eigenvalues $\widehat{a}_l$ are consistent for the $a_l$ (by continuity). The p-value $\mathbb{P}\big(\sum_{l\in L} \widehat{a}_l\, \chi^2_{1,l} > \widehat{S}_n\big)$ is thus asymptotically valid and is computed with a saddlepoint approximation (Kuonen, 1999).

### S2 Simulation details

#### S2.1 GSEA permutations

Gene Set Enrichment Analysis (GSEA) (Subramanian et al., 2005) is a competitive gene set testing method that ranks genes according to a gene-level statistic and computes an enrichment score for each gene set. The significance of the enrichment score can be assessed using either gene-set permutations or label permutations. In the case of gene-set permutation, the null hypothesis is that the gene set tested is not more enriched than a random gene sets. The label permutation tests if the enrichment of the gene set is associated with the variable of interest.

In the R package `fgsea` the function `fgseaLabel` performs GSEA with the classical label permutation. However, in our setting, conditional permutations are required to account for the covariate $\boldsymbol{Z}$. Since the function `fgseaLabel` does not allow for conditional permutations, we did not include GSEA with label permutation in our simulation study. We only included gene permutation, performed with the function `fgseaMultilevel`, with 1000 permutations. We used the default enrichment score as in the original GSEA, and selected the option `pos` to focus on positive enrichment.

#### S2.2 Generation of the gene expression matrix $Y$

In the simulation setting the number of genes and cells are set to $m = 1000$ and $n = 200$, respectively. The covariate $Z \in \mathbb{R}^n$ is generated according to a Gaussian distribution $Z \sim \mathcal{N}(0, 0.1^2)$. The variable of interest $X \in \mathbb{R}^n$ is generated conditionally to $Z$ according a Gaussian distribution $X \sim Z + \mathcal{N}(0, 0.4^2)$. The parameter $\beta = \mathbb{1}_{\{DE\}} \in \mathbb{R}^m$ encodes whether each gene is truly DE or not.

We compared two DE scenarios: Difference in Mean (DMean) and Difference in Modality (DMod). The first one is the classical approach where there is a difference in mean between the conditions, and the second one where truly DE genes have different numbers of modes across conditions.

The gene expression matrix $Y \in \mathbb{R}^{n\times m}$ is defined as $Y = \text{signal} + \epsilon$, where $\epsilon$ is an equi-correlated Gaussian noise matrix. This matrix is parameterized by $\rho \in \{0, 0.4, 0.8\}$, which is the correlation between any pair of genes. The definition of the `signal` depend of the scenario:

- In the mean-difference case the `signal` is the outer product $X\beta$.
- In the modality-difference case the expression of truly DE genes will change according to the value of $X$ to have a bimodal or an unimodal distribution:

$$\texttt{signal} \sim \begin{cases} \mathcal{N}(2, 0.5^2), & \beta = 0 \\ \mathcal{N}(3, 0.5^2), & \beta = 1 \text{ and } X \leq 0 \\ 0.5\mathcal{N}(2, 0.5^2) + 0.5\mathcal{N}(4, 0.5^2), & \beta = 1 \text{ and } X > 0 \end{cases}$$

#### S2.3 Additional simulation results

In the main manuscript we present the results obtained when the pairwise gene correlation is set to $\rho = 0.4$. Here, we report simulations with two more extreme settings, the first one where the genes are independent ($\rho = 0$), as shown in Figure S1, and the second one with a high level of correlation ($\rho = 0.8$), as shown in Figure S2.

Across all three settings, the conclusions remain consistent: self-contained methods generally have greater power, and `cit_gsa` method successfully detects the DMod scenario.

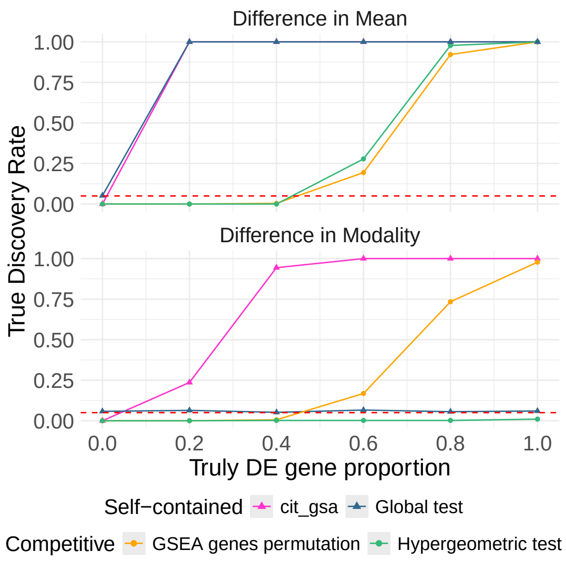


Figure S1: True Discovery Rate as a function of the proportion of truly DE genes in a gene set of size 100, for two DE scenarios: DMean (top) and DMod (bottom) with $\rho = 0$.

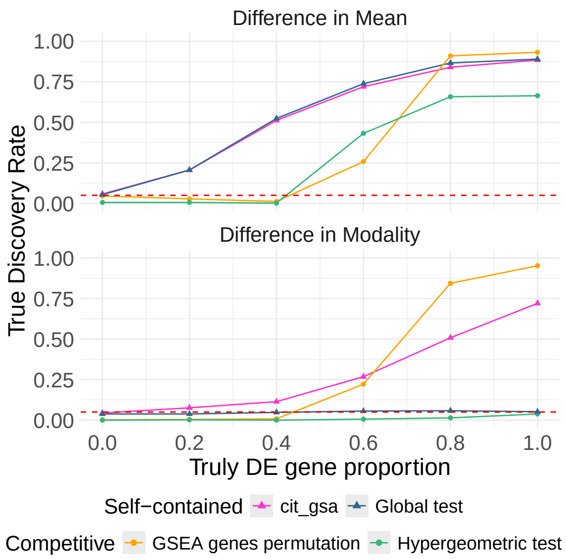


Figure S2: True Discovery Rate as a function of the proportion of truly DE genes in a gene set of size 100, for two DE scenarios: DMean (top) and DMod (bottom) with $\rho = 0.8$.

### S3 Additional numerical results for Villani et al. (2017) data

#### S3.1 Descriptive analysis

Villani et al. (2017) identified marker-genes that are characteristic of the different DC sub-populations: DC1, DC2&DC3 and pDC. The signature highlighted is represented in Figure S3: the expression patterns of these genes indeed allow to separate the cells according to their cellular sub-types.

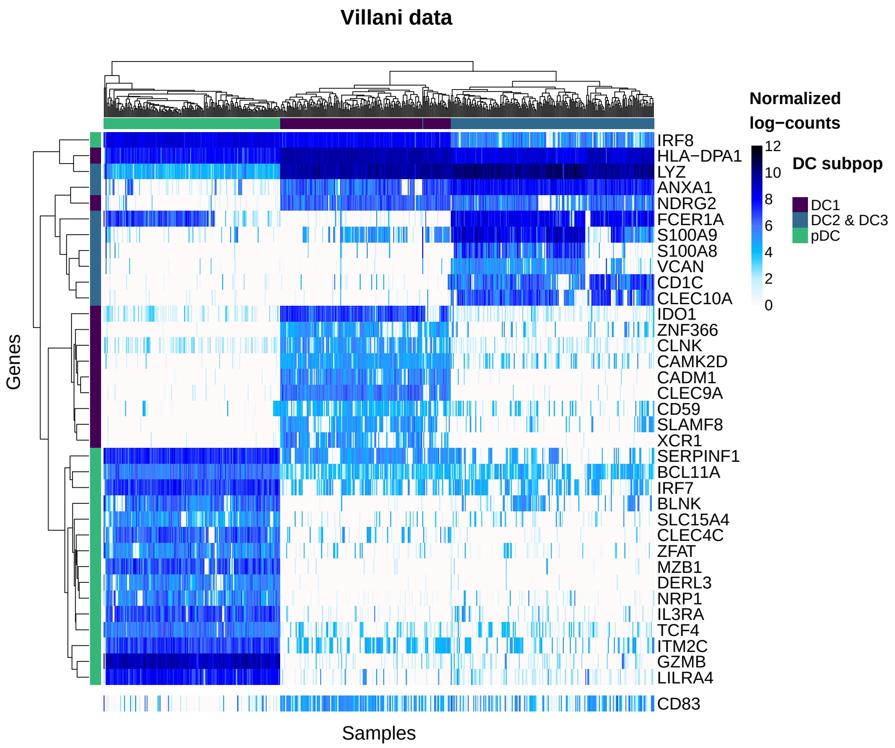


Figure S3: Heatmap of normalized log gene expression of the most characteristic genes of the DC sub-population identified by Villani et al. (2017), including `CD83`.

#### S3.2 CD83 analysis

As described in the main manuscript, we analyzed CD83 gene expression across the three DC subsets. Figure S4 shows that its expression varies across these DC sub-populations, with the *t*-test showing a significant mean difference between all group pairs. Table S1 provides the names of the gene sets that are significant in the conditional gene set analysis. These gene sets are consistent with a state of DC activation.

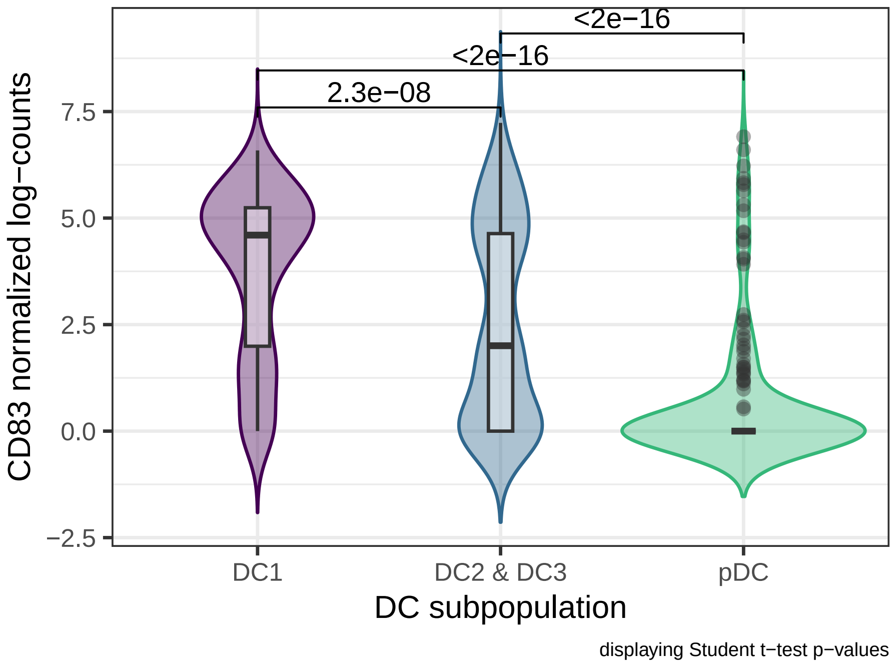


Figure S4: Distribution of CD83 gene expression across the DC sub-populations

| BTM biological annotation |
|---|
| AP-1 transcription factor network |
| cell activation (IL15, IL23, TNF) |
| signaling in T cells (I) |
| myeloid, dendritic cell activation via NFkB (I) |
| putative targets of PAX3 |
| putative targets of PAX3 |
| growth factor induced, enriched in nuclear receptor subfamily 4 |
| enriched in activated dendritic cells (I) |
| leukocyte differentiation |
| enriched for TF motif TTCNRGNNNNTTC |
| DC surface signature (S5) |
| Naive B cell surface signature (S8) |

Table S1: Significant gene sets in the conditional analysis of the gene CD83.

##### S3.3 IDO1 analysis

We performed a similar analysis for the gene IDO1, which is associated with increased immune tolerance of DCs. The expression of this gene also varies across the three sub-populations (Figure S5). But the t-test shows here that there is no significant difference of IDO1 mean expression between the DC2 & DC3 and the pDC. The results of the gene set analysis, presented in Figure S6, shows that among the 259 BTM gene sets of Stanford, one was significant only in the conditional analysis, while four were significant in both the conditional and marginal analyses. In comparison, 238 gene sets were significant only in the marginal analysis. The one gene set identified as significant in the conditional analysis only is indicated by an asterisk in Table S2.

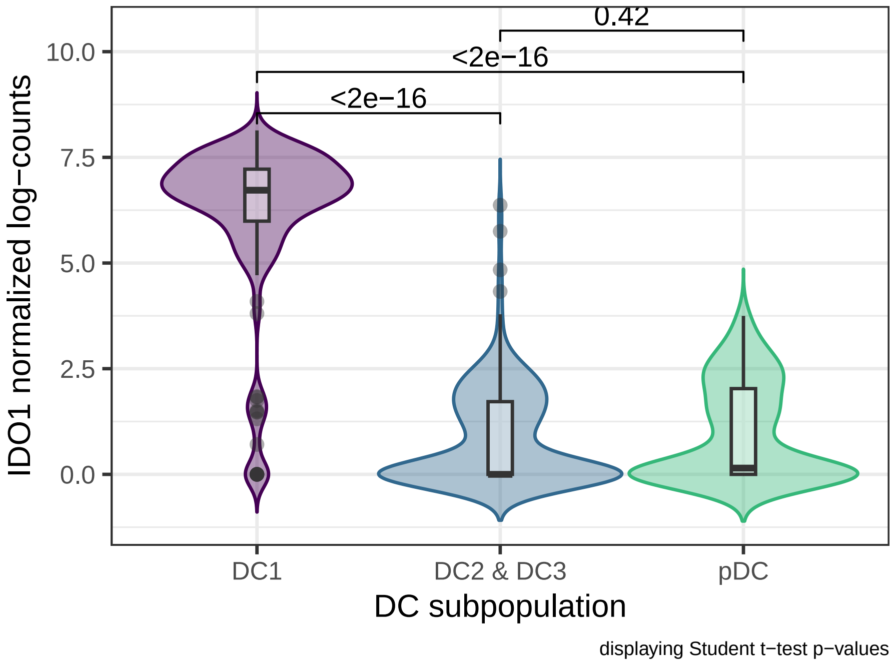


Figure S5: Distribution of IDO1 gene expression across the DC sub-populations.

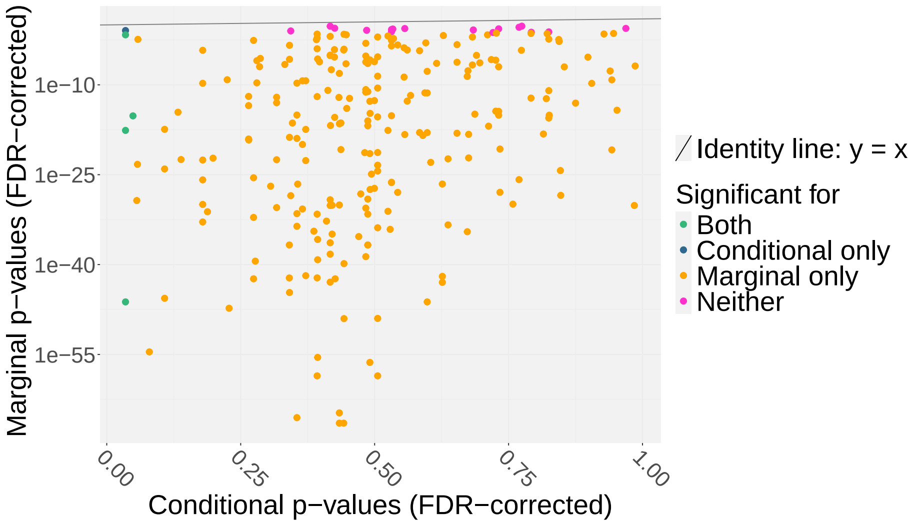


Figure S6: Gene set analysis results for the association between IDO1 expression, comparing for each of the 259 BTM gene sets of Li et al. (2014) the conditional $p$-value after DC subset adjustment ($x$-axis) with the marginal $p$-value with no such adjustment ($y$-axis). The FDR-corrected $p$-values are shown on a logarithmic scale, with a break in the $x$-axis. The solid line represents the identity line.

| **BTM biological annotation** |
|---|
| * signal transduction, plasma membrane |
| integrin cell surface interactions (I) |
| extracellular matrix (I) |
| chemokines and inflammatory molecules in myeloid cells |
| enriched in extracellular matrix & associated proteins |

Table S2: Significant gene sets in the conditional analysis of the gene IDO1. The gene set ”signal transduction, plasma membrane” is significant only in the conditional analysis.
